# BinaRena: a dedicated interactive platform for human-guided exploration and binning of metagenomes

**DOI:** 10.1101/2022.06.28.498025

**Authors:** Michael J. Pavia, Abhinav Chede, Zijun Wu, Hinsby Cadillo-Quiroz, Qiyun Zhu

## Abstract

Exploring metagenomic contigs and “binning” them are essential for delineating functional and evolutionary guilds within microbial communities. Despite available automated binners, researchers often find human involvement necessary to achieve representative results. We present BinaRena, an interactive graphic interface dedicated to aiding human operators to explore contigs via customizable visualization and to associate them with bins based on various data types, including sequence metrics, coverage profiles, taxonomic assignments and functional annotations. Binning plans can be edited, inspected and compared visually or using algorithms. Completeness and redundancy of user-selected contigs can be calculated real-time. We show that BinaRena facilitated biological pattern discovery, hypothesis generation and bin refinement in a tropical peatland metagenome. It enabled isolation of pathogenic genomes within closely-related populations from human gut samples. It significantly improved overall binning quality after curation using a simulated marine dataset. BinaRena is an installation-free, client-end web application for researchers of all levels.

## Introduction

The rapid advancement in high-throughput sequencing technologies has led to the discovery of an enormous amount of new biodiversity from uncultivated microbial populations ^1^. Extracting population genomes from heterogenous microbial communities is essential to understand the contribution of defined microbial lineages to host and environmental processes. Genome-resolved metagenomic studies have provided valuable insight into understanding microbial links to biogeochemistry ^2–5^, connections to human health and disease ^6–8^, and discovery of novel microbial groups ^9,10^. Exploration of such datasets can quickly become cumbersome ^11–15^, comprising hundreds of metagenome-assembled genomes (MAGs) with associated sequence characteristics, functional potential, and abundance across samples.

The building block of this comprehensive data can be reduced into a contig, the minimum unit of a genomic sequence derived from the assembly of metagenomic reads. Using characteristics such as nucleotide composition and sequencing depth, similar contigs can be grouped into “bins” representative of microbial populations’ genomes (i.e., MAGs). Despite the wealth of automatic binning tools ^16–19^, an intermediate step which can contextualize multiple layers of user-specified information for inspection of contig-to-bin assignment is necessary for reliable conclusions to be made. This human-guided step can greatly improve the quality of bins and subsequent inferences made from the contained biological information ^1,20–22^. This is because human brains are highly effective in pattern recognition ^23^, which was only recently challenged by algorithms in limited tasks ^24^, and this ability can be further enhanced by a priori knowledge of the biological systems. It has been accepted that exploratory data analysis ^25^, as characterized by heavy employment of data visualization and human involvement, is essential for understanding complex datasets, removing noises, discovering patterns, and generating hypotheses ^26^, and this cannot be replaced by any uniform algorithmic workflow.

Therefore, software infrastructure that helps human researchers in exploring metagenomic assemblies and defining bins (MAGs) is much needed ^27^. Multiple tools have been developed to provide interactive visualization of metagenomes ^28–32^ (reviewed below), which can facilitate this process. However, few are explicitly designed with the goal of maximizing human productivity. Most tools constrain usability either through computational skill thresholds, or a relatively inflexible workflow, or a lack of study specific customizable features, respectively.

To address this gap, we present BinaRena (“bin arena”), a comprehensive, highly customizable interactive graphical interface dedicated to human-guided exploration and binning of metagenomes. A visual representation of contigs are rendered as a scatter plot, displaying flexible types of data such as sequence metrics, coverage profiles, *k*-mer frequency, taxonomic assignment, feature annotation, existing binning outputs, and other metrics appropriate to the researcher. Integration of multiple layers of contig characteristics can aid delineation of microbial community members and improve overall binning results. The BinaRena program is free of installation, dependency, and a web server, making it exceptionally convenient for deployment and use. Licensed under BSD-3-Clause, BinaRena’s source code is hosted at: https://github.com/qiyunlab/binarena, together with documentation, example data, and a fully functional live demo.

To demonstrate BinaRena’s functionality and how it improves microbiome research, we analyzed one synthetic and two real-world metagenomic study cases. Specifically, we (1) analyzed the first metagenome available of a complex open tropical peatland from Maquia (MAQ) within the Pastaza-Marañón Foreland Basin, a globally important carbon reservoir in the Amazon, (2) re-analyzed metagenomes confounded by multiple pathogens from fecal samples of Traveler’s Diarrhea (TD) patients ^33^, and (3) quantified the systematic improvement of binning results using the gold standard CAMI2 marine dataset ^19^. We show that BinaRena significantly facilitated pattern discovery, hypothesis generation, strain-level isolation and bin refinement that were otherwise not achievable or overlooked by automatic workflows.

## Results

### Design and functionality of BinaRena

BinaRena is an installation-free, client-end web application. The user may simply double click “BinaRena.html” in the downloaded package to launch the program, which is literally a single webpage running in the user’s web browser, and does not require a web server running in the backend. In this sense, it is analogous to bioinformatics programs like Krona ^34^ and EMPeror ^35^. However, BinaRena further removes the need for executing a script to construct the webpage. Instead, the user may simply drag and drop data files into the browser window to load them. This design minimizes the efforts for deployment and preparation, especially for non-technical users, and computer systems with restrictions. The BinaRena program is written in pure JavaScript, without using any third-party frameworks or libraries. This ensures the program’s flexibility in behavior and functionality, and allows the developing team to optimize the code for improved performance in rendering and calculation, which is important for handling modern metagenomic datasets, which usually contain tens to hundreds of thousands of contigs and many properties.

The main workspace of BinaRena is an interactive scatter plot, with data points representing contigs from an assembly (Fig. 1). The plot appearance is defined by five aesthetics: *x*- and *y*-axes, size, opacity, and color, each of which can be customized in the interface based on user-provided data that are relevant in delineating or relating contigs. For example, plotting GC% by coverage, comparing per-sample abundance profiles, and *k*-mer frequency-based dimensionality reduction, are all helpful for contig clustering ^28,32,36^. Reference-based properties such as taxonomic assignment and functional annotation further inform the biology of contigs. BinaRena enables convenient toggling among these characteristics. The user may further specify data transformation, data range, and color map (for both discrete and continuous data) in the interface. The program implements multiple transformation methods to deal with various types and distributions of biological data, including square and cube (root), logarithmic and exponential, logit and arcsine, and ranking, all of which can be easily triggered from a dropdown menu. The user may move and zoom the plot with mouse and/or keyboard, just like navigating a typical digital map. All panels can be uncollapsed to screen corners to minimize distraction during data exploration. The scatter plot along with legends and axes can be exported as a PNG bitmap image or an SVG vector image for post-processing and publication.

**Figure 1.**
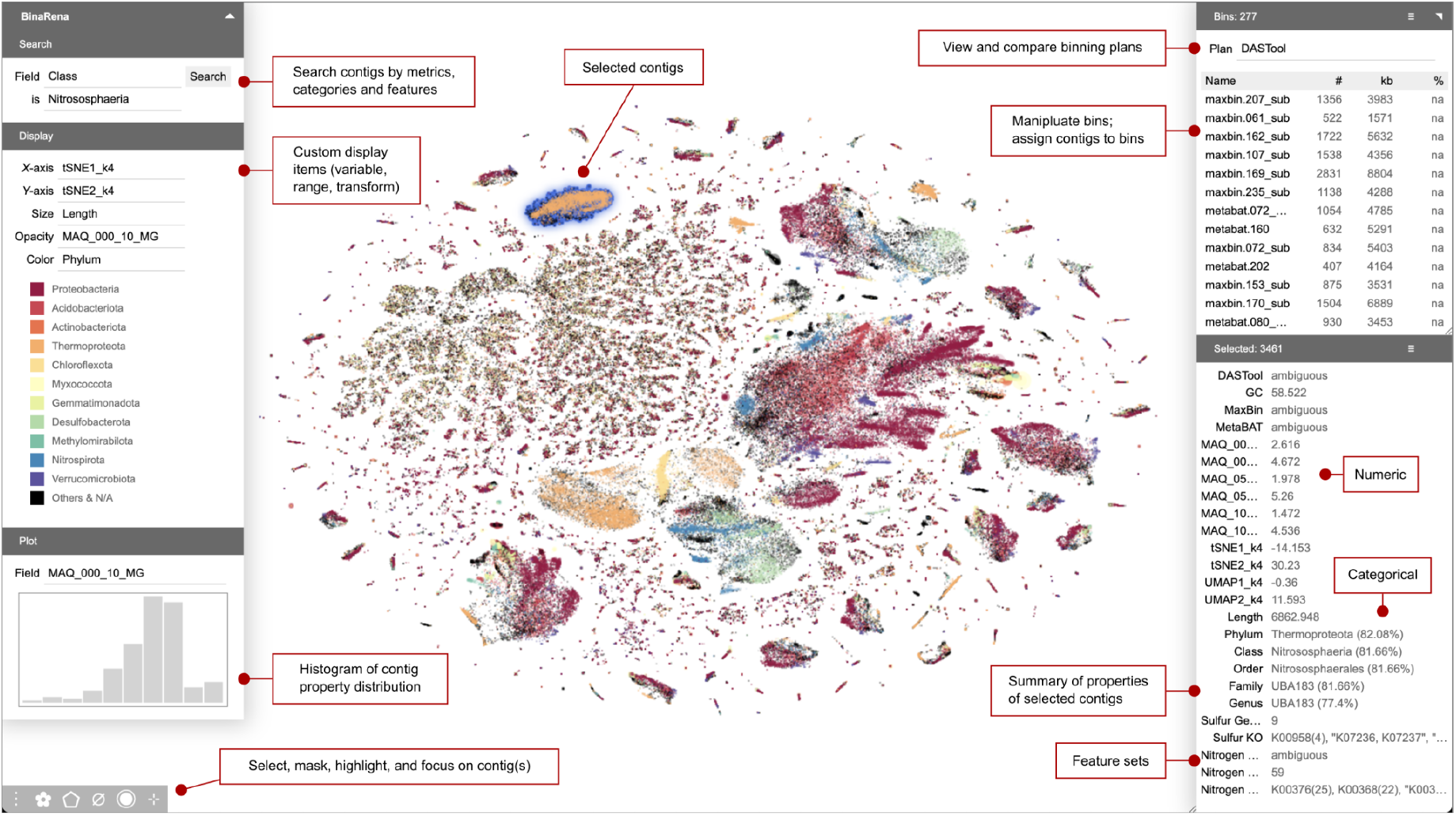
A screenshot of the main interface of BinaRena. The program is displaying the MAQ dataset, consisting of 262,705 contigs obtained from a co-assembly of six tropical peatland metagenomes. *X*- and *y*-axes represent t-SNE embeddings based on tetranucleotide frequencies. Marker size (radius) is proportional to the cube root of contig length. Marker opacity is proportional to the logarithm of sequencing depth (coverage) in one sample. Colors are assigned to the 10 most frequent phyla binned from DASTool. A binning plan consisting of 277 bins pre-computed by DASTool is loaded in the program, allowing the user to explore and manipulate individual bins by adding / removing individual contigs. A spatially distinct cluster of 3,461 contigs putatively representing multiple Nitrososphaeria MAGs is currently selected by the user. The properties of the selected contigs are summarized in a side panel. The distribution of coverage is displayed as a histogram. Red-edged text boxes indicate functional components of the BinaRena interface.

The contig data can be provided as one integrated data table, or as multiple tables or mappings sequentially appended to the same dataset, which improves flexibility and lowers the challenge in preparing input files. BinaRena accepts four data types: numeric, categorical, feature set and descriptive. The feature set data type, provided as comma-separated strings, lets the user specify gene content of each contig, annotated either for general purpose (such as KEGG ^37^ Ontology, or KO) or to address specific research questions (such as phylogenetic markers ^38^, antimicrobial resistance genes, mobile genetic elements, or members of a specific metabolic pathway).

BinaRena further lets the user specify feature groups, defined by a list of member features that constitute a group. Then the program can calculate the completeness and redundancy (a.k.a., contamination) of user-selected contig groups in real-time. This significantly improves the convenience and flexibility to assess the quality of a putative bin with or without a specific biological question, as in contrast to currently adopted protocols which are usually performed when bins are already defined. It should be noted however, that BinaRena does not consider marker gene collocation as CheckM does ^38^, therefore their results are not identical, albeit highly correlated (Fig. S1), thus the former can serve as a first-pass check while the latter is still recommended post binning.

BinaRena offers a variety of controls for exploring the metagenomic dataset. Contigs can be selected by mouse clicking, or by drawing a polygon to contain multiple contigs. The selection retains as the aesthetics are toggled, allowing the user to explore the same contigs of interest using different data. With a single keystroke or button, the selected contigs can be highlighted using choice of colors to indicate user interest; they can be “masked” such that they are both hidden from the plot and excluded from subsequent manipulations and calculations; they can be “focused” such that only them but no other contigs are visible, which facilitates user concentration. These operations can be “undone” to revert to previous status. Contigs can be searched based on their numeric and categorical properties as well as features they carry.

The properties of selected contigs are summarized in a side panel by user-specified methods that make most sense for the nature of data. Examples are: “Length” is the sum of contig lengths. “GC” is the average of GC contents weighted by length. The category (such as taxonomic group) of multiple contigs is determined by the majority rule, optionally weighted by length, with the fraction annotated as a prefix (e.g., “Firmicutes (80%)”). Aside from the scatter plot, there is a mini interactive histogram displaying the distribution of a user-designated numeric property (such as coverage) of the selected contigs. The user can use mouse dragging to filter the contigs by data range (such as a peak of coverage values). This function is useful for refining a contaminated bin.

BinaRena provides handy controls for assigning contigs to bins that represent putative MAGs. The user can create a binning plan *de novo* or edit binning plans computed by external programs. Bins are displayed in an interactive table and summarized by their total length and abundance per sample. Using one keystroke or button, the user can add or remove selected contigs to or from individual bins. The binning plan or the contig data of individual bins can be exported to table files. BinaRena implements two algorithms for the evaluation and comparison of binning plans.

It calculates the silhouette coefficient ^39^ to assess the confidence of assigning contigs to individual bins. The results can be visualized instantly as color depth to provide an intuitive view of the bin confidence profile. The program also calculates the adjusted Rand index ^39^ to assess the consistency between pairs of binning plans. Both metrics are widely used in cluster analysis. However BinaRena’s ability to calculate them during exploration significantly supports the user effort.

Besides the main program, BinaRena provides multiple Python scripts to aid data preparation. They include utilities to count *k*-mers from contig sequences, then to conduct dimensionality reduction using PCA, t-SNE and UMAP, and utilities to convert common metagenomics tool outputs into the table format. Examples are SPAdes ^40^ and MetaHIT ^41^ assemblies, GTDB-tk ^42^ lineage strings, Kraken ^43^ taxonomic assignments, and CheckM ^38^ marker gene maps.

A video introduction to BinaRena’s functionality is provided in Data S1.

### Comparison with existing tools

Here we review multiple existing tools for interactive visualization of metagenomic contigs and compare them with BinaRena. Anvi’o ^28^ is an integrated multi-omics platform that is most known for an interactive sector graph depicting sequence composition and per-sample abundance of contigs with the ability to add customizable layers, allowing users to explore classification, evolutionary and functional capacity patterns of the dataset. This visualization method is highly effective for exploring contig distribution among samples, but less so for the relationships among contigs. The complexity in setting up a server and executing command-line workflows to prepare for visualization may challenge non-technical users. The visualization tool ICoVeR ^29^ is for user-guided refinement of existing binning plans. It renders a line graph depicting per-sample abundance of a co-assembly, as well as other numeric metrics. It supports generation of scatter plots and histograms using several clustering and ordination algorithms, however these plots are for exploring variables instead of contigs. The ggKBase ^30^ workflow is suitable for manual binning. It employs an interactive wheel for selecting taxonomic groups and a histogram for selecting metric ranges (also supported by BinaRena). Collectively, BinaRena’s interactive scatter plot of contigs does not overlap with Anvi’o, ICoVeR and ggKBase, but instead may serve as a complement to current metagenomics workflows that use these tools. BusyBee Web ^31^ is a web server that performs the entire binning workflow. Its interface displays contigs as a scatter plot, which is mainly for exploring pre-computed (by the server) bins, and does not support complex contig and bin operations. Likewise, it displays CheckM-calculated bin quality metrics, rather than evaluates bin quality interactively as BinaRena does. We would like to note that BusyBee Web’s predecessor, VizBin ^36^, was the original source of inspiration to the development of BinaRena. To our knowledge, Elviz ^32^ is the most comparable existing tool to BinaRena. The Elviz server is integrated into the JGI portal, which provides convenience but also imposes restrictions to the user. It emphasizes on assessing the taxonomy and functions of contigs, but it can be repurposed for editing bins. An itemized comparison of BinaRena and Elviz is provided in Table S1, showing that the former is notably more feature-rich. Finally, an obvious advantage of BinaRena compared with all these tools is the ease of deployment. In summary, we believe that BinaRena is a unique bioinformatics tool for the task it aims to achieve.

### Exploring microbial populations responsible for nutrient cycling in the Maquia peatland

Extensive tropical peatland formations have been reported in the Amazon ^44^, of which the “open peatland” is a unique category devoid of trees but dominated by arbustive vegetation and widespread in the Pastaza Marañon basin ^45^. Given their role sequestering organic carbon in their soils and to understand their microbial functions ^46^, we sampled an open peatland (Maquia: MAQ) for metagenomic evaluation. Quality-filtered reads from all six samples were co-assembled and subsequent contigs were binned using three automatic binners: MaxBin ^16^, MetaBAT ^17^, and DASTool ^18^, yielding 251, 345, and 276 total bins, respectively, spanning 25 phyla. BinaRena was used to render placement of contigs from this assembly (contigs ≥ 2000 bp) based on t-SNE on tetranucleotide frequency (Fig. 1). The contig aesthetics (size, color, and opacity) are associated with common properties such as contig length, taxonomic classification, and abundance in a sample from depth = 10cm (see Methods). This initial view revealed that many contigs are associated with taxonomic groups from Proteobacteria, Acidobacteria, and Actinobacteria. Additionally, Methylomirabilota, Desulfobacterota, and Nitrospirota form two distinct tight clusters. BinaRena’s polygon tool was used to select a distinct cluster of contigs, representing multiple populations of Nitrososphaeria. Classified under phylum Thermoproteota, Nitrososphaeria are ubiquitous terrestrial ammonia oxidizing archaea ^47^.

Genome-resolved metagenomics is widely used to understand potential biological mechanisms within an ecosystem and its distribution across the community. Nitrogen cycling within tropical peatlands is relatively understudied, yet there is evidence that it is closely interconnected with release of greenhouse gasses from these environments ^48,49^. Here we used BinaRena to explore the distribution of nitrogen cycling genes involved in pathways such as dissimilatory nitrate reduction, denitrification, nitrification, and nitrogen fixation. Visualization in BinaRena supports quick identification of contigs containing genes for the previously listed pathways of interest (Fig. 2A). The distinct cluster of Nitrosophaeria contigs contain copies of both the *nosZ* and *nirK* (Cluster 1). Additionally, a different cluster of Nitrosophaeria contigs (Cluster 2) was found with copies of *amoABC*, which is consistent with culture-based studies ^50^ of Nitrososphaeria as an ammonia oxidizer. Overall, BinaRena assisted in identification and quantification of the importance Nitrosophaeria potentially carry out in the MAQ nitrogen cycle.

**Figure 2.**
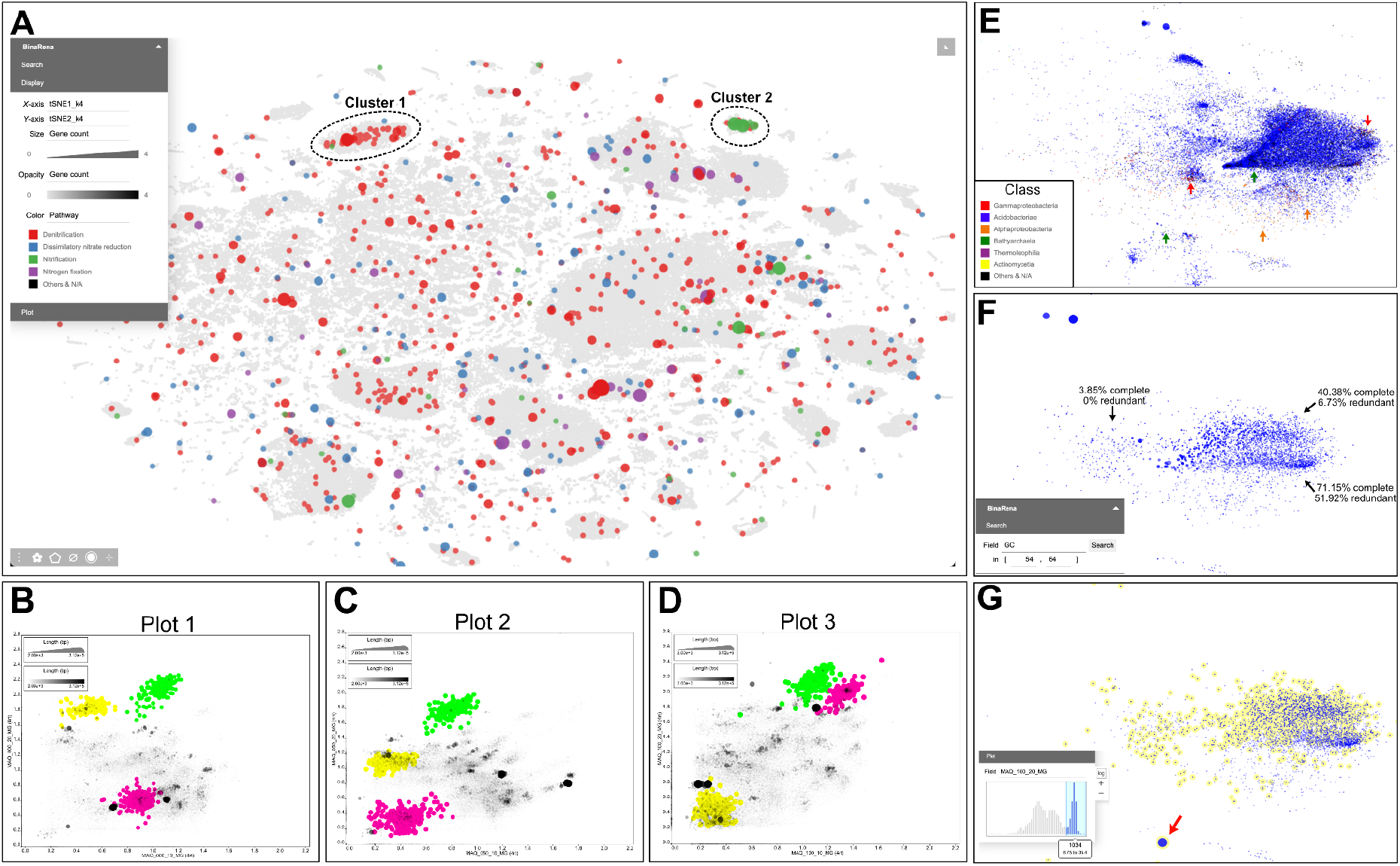
Distribution of nitrogen cycling genes and exploration of *Sulfotelmatobacter* populations in the MAQ dataset. **A**. An overview of the entire assembly. The *x*- and *y*-axis represent t-SNE embeddings based on tetranucleotide frequencies. Marker size (square root) and opacity is proportional to the number of KOs assigned to each contig that are associated with the previously described nitrogen pathways and the color represents that pathway. **B**-**D**. BinaRena-exported SVG images (rasterized) depicting the change in abundance of the Streptosporangiales MAG (pink), Thermoanaerobaculales MAG (yellow), and Nitrososphaerales (green). The only edits to raw files generated by BinaRena were an increase in font size, changes to legend text, and resizing of plot area to decrease white space. **E**. Subset of contigs classified as Koribacteraceae or were assigned to one of the five *Sulfotelmatobacter* MAGs are plotted using t-SNE (*k*=6), and colored by class. The size is proportional to the contig length and opacity is the cube for coverage in location 3, depth 20cm. Arrows are pointing at regions with contigs of potential contamination. **F**. Contigs in panel E that have been filtered to a range of 54-64% GC (inset). All other aesthetics remain the same. **G**. Contigs highlighted in yellow were selected based on high abundance in location 3, depth 20cm using the histogram (inset). All other aesthetics remain the same except for size which is proportional to cube root of the amount of sulfur genes found on contigs. The red arrow is pointing at the potentially missing contig from the *Sulfotelmatobacter* MAG.

To understand how there might be differences in nitrogen cycling populations across the 100m transect, we focused on two high- and one medium-quality bins (defined following ^1^) inferred by DASTool that were identified as capable of dissimilatory nitrate reduction (Streptosporangiales: 96.77% complete / 3.83% redundant, calculated by CheckM, same below), and denitrification (Nitrososphaerales: 76.79% / 0.93% and Thermoanaerobaculales: 94.12% / 4.2%) (Fig. 2B-D). Contigs in the Thermoanaerobaculales MAG form a distinct cluster in the BinaRena graph representing the coverage profile at both 10cm and 20cm depths at location 1(Fig. 2B). This MAG is predicted to carry out nitrite reduction (*nirK*), a suboxic process ^51^, and likely why we find it at a higher abundance at 20cm in the soil. However, abundance of this MAG progressively decreases in location 2 and then location 3 (17.92× to 4.41× to 0.07×). Conversely, the Streptosporangiales MAG is found at very low abundance at both locations 1 and 2, but becomes abundant at 10cm (5.2×) and 20cm (14.12×) depths in location 3. While we observed spatial variation in the abundance of both the Thermoanerobaculales and Streptosporangiales MAGs there was minimal variation detected in Nitrososphaerales. The Nitrososphaerales was the most abundant MAG across all three locations (31.83×, 33.94×, 18.79×) at 20cm depth. It is interesting to consider what environmental factors are contributing to the change in abundance of both the Thermoanaerobaculales and Streptosporangiales MAGs. However, this falls outside the scope of this study, but demonstrates BinaRena’s utility in hypothesis generation.

BinaRena is capable of restructuring contig placement expediting identification of dynamics between populations while also supporting MAG refinement by identifying potentially misplaced contigs (and genetic potential). The recently discovered genus of *Sulfotelmatobacter* is potentially capable of carrying out dissimilatory sulfite or sulfate respiration ^52^, with implications for organic matter decomposition and greenhouse gas production. The *Sulfotelmatobacter* MAGs recovered using the three automated binners were relatively low quality (high redundancy or low completeness), as well as, lacked genes involved in sulfur metabolism (Fig. S2). To improve MAG quality we selected all contigs associated with these five MAGs, and contigs that were classified as Koribacteraceae and then subsequently visualized using the t-SNE at *k* = 6 (Fig. 3E). We found most contamination (from contig-to-MAG selection) coming from contigs classified as Alphaproteobacteria (280), Gammaproteobacteria (61), and Bathyarchaeia (15), which were selected and removed. After removal, we re-assessed the distribution of contigs and selected those that fell within a tight range of GC content (54-62%, based on MAGs generated from the automatic binners, as well as, what has been previously published on this group ^52^) and were removed using BinaRena.

**Figure 3.**
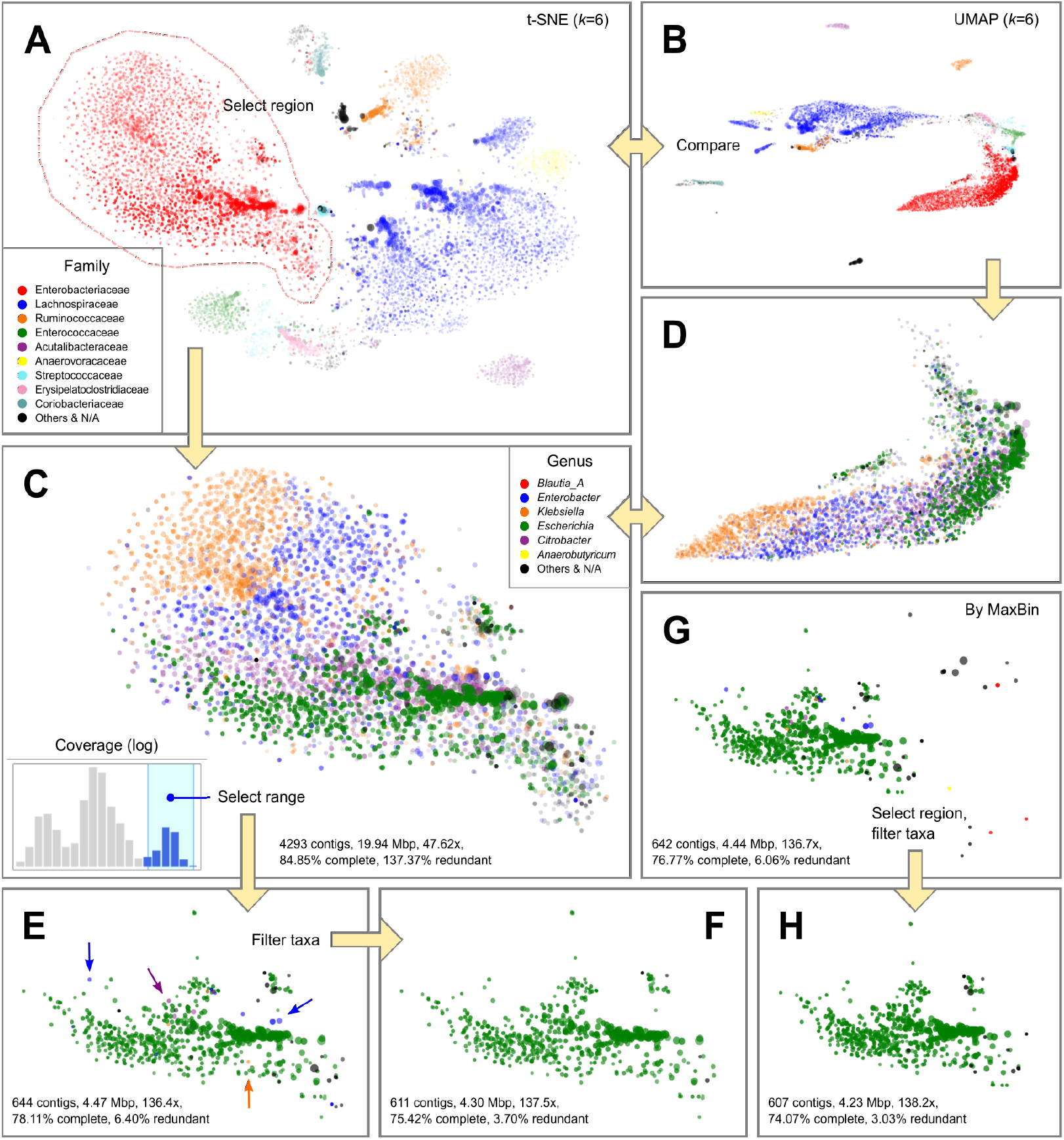
Extraction of a pathogenic *Escherichia coli* MAG from several closely related organisms in the metagenome from the gut of a Travelers’ Diarrhea patient (sample #76, with 2.92 Gbp raw reads, 10,910 contigs totaling 69.5 Mbp). Marker size (radius) is proportional to the cube root of contig length. Marker opacity is proportional to the cube root of contig coverage. Colors were assigned to the most abundant taxa in the sample. The assembly data (**A**, **B**) was explored using alternative dimensionality reduction methods (t-SNE for **A**, **C**, **E**-**H**; UMAP for **B**, **D**, both based on *k*-mer (*k* = 6) frequencies). A distinct blob of Enterobacteriaceae contigs (A, dashed line) were selected (**C**, **D**), and filtered based on its coverage profile (**C**, inset), resulting in a putative *E. coli* bin (**E**), which was further filtered by taxonomy (exemplified by arrows in **E**) to improve purity (**F**). In parallel, the corresponding bin inferred by MaxBin (G) was filtered by spatial pattern and taxonomy to retain a purer bin (**H**).

There were three visually distinct clusters of contigs, but binning these resulted in either low completeness or high redundancy (worse than the automated binners) (Fig. 3F). To better account for differences between populations we further focused on contig abundance across location and depth. Using BinaRena’s interactive histogram we separated contigs that were at high abundance in location 3 at 20cm depth (*Sulfotelmatobacter* are predicted anaerobes, and the MAGs from the automated binners were the most abundant in location 3 at 20cm) (Fig. 3G). This retained 1,034 contigs with a total size of 3.98 Mbp and an average of 13.95× coverage. Further the completeness and redundancy of this MAG marginally increased to 58.55% and 3.41%, respectively. This MAG was comprised of 78.66% of the MetaBAT #229 bin and 99.4% of the Maxbin #235_sub bin from DASTool. While quality only slightly increased the MAG now contained genes for dissimilatory sulfate respiration (*dsrAB*) (Fig. 3G, red arrow). These genes were previously found by MaxBin (#235), but were removed by DASTool. To support their placement within this bin, both *dsrA* and *dsrB* genes were blasted and the top ten matches belong to an uncultured sulfate-reducing organism. This 2,139 bp-long contig does not cluster with the rest of the contigs (based on t-SNE k=6). We suggest that given contigs ≤ 2000 bp are known to have issues binning ^21^ it is possible that this contig was misplaced by the automated binners. By implementing both targeted classification, GC% and depth metrics (for what is known about *Sulfotelmatobacter*) we were able to recover a more complete representation of the ecosystem. In summary, BinaRena directly facilitated the curation of this MAG, which prior to human intervention lacked biological significance.

### Separating closely-related pathogenic microbes in Traveler’s Diarrhea gut metagenomes

Travelers’ Diarrhea (TD) is an intestinal disorder caused by infection during traveling ^53^. Identification of infectious agents is of epidemiological importance but challenging due to the diverse and unpredictable pathogenic profiles ^54^. In a previous study, Zhu et al. studied the metagenomes of a TD cohort, and discovered multiple putative pathogens, some of which were confounded by closely related organisms in the same sample ^33^. The current study provides a re-visit to the question using the BinaRena program, as exemplified by two difficult samples.

Sample #76 was characterized by the co-infection of multiple putative pathogens under the genera of *Escherichia, Enterobacter*, *Klebsiella*, and *Citrobacter*, all belonging to the family of *Enterobacteriaceae* ^33^. The evolutionary proximity of these organisms increases the challenge of assigning contigs to correct genomes. The relatively shallow sequencing depth (2.92 Gbp raw reads in total) further adds to the difficulty in recovering MAGs of reasonable quality. We performed visual observation of the assembly in BinaRena, showing that t-SNE and UMAP at *k* = 6 provided the most apparent visual consistency between contig clustering pattern and taxonomic assignment (Figs. 3A, B, S3). By cross-comparing the two views, we selected a cluster of contigs that are mainly assigned to *Enterobacteriaceae* using BinaRena’s polygon selection tool. BinaRena reported that this cluster contains 4,293 contigs totaling 19.9 Mbp, with an average coverage of 47.62× (weighted by contig length). 98.42% of the length was assigned to family *Enterobacteriaceae*. By assessing CheckM’s *Enterobacteriaceae*-specific marker genes (*n* = 297), BinaRena determined that this cluster has completeness = 84.85% and redundancy = 137.37%, indicating the presence of multiple genomes (Table S2, same below). Coloring by genus clearly showed that this cluster contains contigs assigned to all four pathogenic genera, which are visually distinguishable but hard to separate (Figs. 3C, D). The histogram of contig coverage showed several peaks, again implicating the presence of multiple genomes (Fig. 3E, inset). We separated the high-end peak by mouse-dragging six bins out of 20 in the interactive histogram (Fig. 3E, inset). This retained 644 contigs (4.47 Mbp, 136.4×), with 95.87% of its length assigned to genus *Escherichia* (Fig. 3E). They are 78.11% complete and 6.40% redundant. Next, we used BinaRena’s search tool to identify and remove non-Gammaproteobacteria contigs, and contigs assigned to the other three pathogenic genera (*Enterobacter*, *Klebsiella*, and *Citrobacter*), which are presumably contaminations. This left 611 contigs (4.30 Mbp, 137.5×), with completeness = 75.42% and redundancy = 3.70%, which we consider as a putative MAG of *Escherichia* (Fig. 3F), a taxon containing common causative pathogens for TD ^54^.

We then explored binning plans generated by automatic binners (MaxBin, MetaBAT, and DASTool). BinaRena’s information panel indicates that MaxBin’s bin #001 (642 contigs, 4.44 Mbp, 136.7×, 76.77% complete, 6.06% redundant) has the highest consistency with the manually isolated *Escherichia* MAG as detailed above (97.08% of the latter length was shared between the two; Jaccard index = 0.901) (Fig. 3G). However, this bin contains multiple “outlier” contigs that are approximate to other clusters indicated by both *k*-mer signature and taxonomic assignment, implicating contaminations (Fig. S4A). Therefore, we manually refined this bin by removing the “outlier” contigs using polygon, then by taxonomic filtering as detailed above (Fig. 3H). The curated bin has 607 contigs (4.23 Mbp, 138.2×), with completeness = 74.07% and redundancy = 3.03%, and has higher consistency with the manually extracted MAG (Jaccard index = 0.952). In parallel, MetaBAT recovered a bin (403 contigs, 88.89% complete, 30.98% redundant) which is a mixture of a portion of *Escherichia* contigs and a clearly separate cluster of contigs that were assigned to genus *Faecalibacterium*, a common commensal component of the gut microbiota ^55^. This observation points to putative chimerism (Fig. S4B). Finally, the ensemble method DASTool kept the MetaBAT bin, and stripped the shared part from the MaxBin bin, leaving only 304 contigs (39.06% complete, 2.36% redundant) (Fig. S4C), a result that is suboptimal.

In parallel, we investigated sample #50076, characterized by the co-infection of multiple *Escherichia coli* strains ^33^. An overview of the assembly in BinaRena supports a clear *E. coli* dominance pattern (Fig. S5A). Among the 27 bins inferred by MaxBin, seven have more than 50% of their total length assigned to genus *Escherichia*, however four of them are less than 2% complete as evaluated by BinaRena using CheckM’s *E. coli*-specific marker genes (*n* = 1,628). The remaining three have a total length between 1.2 and 1.7 Mbp, average coverage between 2100 and 3100×, completeness between 18 and 41%, and redundancy below 0.5% (Table S3). These metrics indicate that they are highly incomplete *E. coli* genomes. The relatively even coverage values and the *a priori* knowledge that *E. coli* genomes are usually 4.5-5.5 Mbp long ^56^ led us to postulate that these bins may be parts of one *E. coli* genome. Similarly, MetaBAT inferred two *E. coli* bins (one was retained by DASTool), each of which also seemingly partial (Table S3). These results expose the limitation of automatic methods in recovering complete pathogen genomes. Therefore, we resorted to *de novo* binning using BinaRena. Similar to the method described above, we first selected the cluster of contigs that were dominantly assigned to *Escherichia*, with 554 contigs, 6.89 Mbp, 1939×, 98.40% complete, and 4.91% redundant (Fig. S5B). These metrics indicate that there may be secondary *E. coli* genomes mixed in it, which is also evident from the multi-modal pattern of the contig coverage histogram (Fig. S5A, inset). Likewise, we selected the top five bins out of 20 (coverage ≥ 1110×), resulting in 309 contigs, 5.07 Mbp, 2515×, 98.03% complete and 1.29% redundant (Fig. S5C). Compared with the automatically inferred bins, this bin notably better represents a complete *E. coli* genome that dominated the patient’s gut amongst other less abundant *E. coli* strains.

### Systematic improvement of binning results using the CAMI2 marine metagenomes

We further demonstrated that BinaRena can help to efficiently and systematically improve bin quality of an entire dataset. For this purpose we used the synthetic marine metagenomic dataset from the 2nd CAMI Challenge, a gold standard for assessing the performance of metagenome binning algorithms ^19^. A researcher worked on each binning plan generated by MaxBin, MetaBAT and DASTool. Briefly, contigs associated with each bin were highlighted and then the *x*- and *y*-axes were toggled (between both PCA and coverage profiles) to identify potential misplaced contigs. The binning plans pre- and post-curation were evaluated using the silhouette coefficient and the adjusted Rand index (ARI), both calculated in the BinaRena interface, and the completeness and redundancy scores calculated by CheckM (outside BinaRena). It should be noted that the researcher was agnostic about these metrics during curation.

Comparative analysis showed that after curation using BinaRena, the binning plan was visibly more consistent with the clustering pattern of contigs, as indicated by silhouette (Fig. 4A, B). BinaRena’s capability of calculating and visualizing silhouette as the user modifies the binning plan is useful to curation (although not used in this analysis). A portion of contigs were filtered out from the bins during curation (Fig. 4C), accompanied by infrequent deletion of entire bins (Fig. 4D). The removed contigs were usually small, therefore the loss in the total bin length was moderate (Fig. 4E). As evaluated by ARI, the post-curation binning plans are notably more consistent with the ground truth genome assignment, as compared to the pre-curation ones (Fig. 4F). For example, ARI of the DASTool-inferred bins increased from 0.729 to 0.982, suggesting that the latter are a nearly perfect subset of the true genomes. The quality of curated bins following the adopted standard ^1^ suggested a notable increase in the number of all publishable MAG categories (high-, medium- and low-quality) (Fig. 4G). These results indicate a substantial improvement in the overall quality of binning plans using BinaRena.

**Figure 4.**
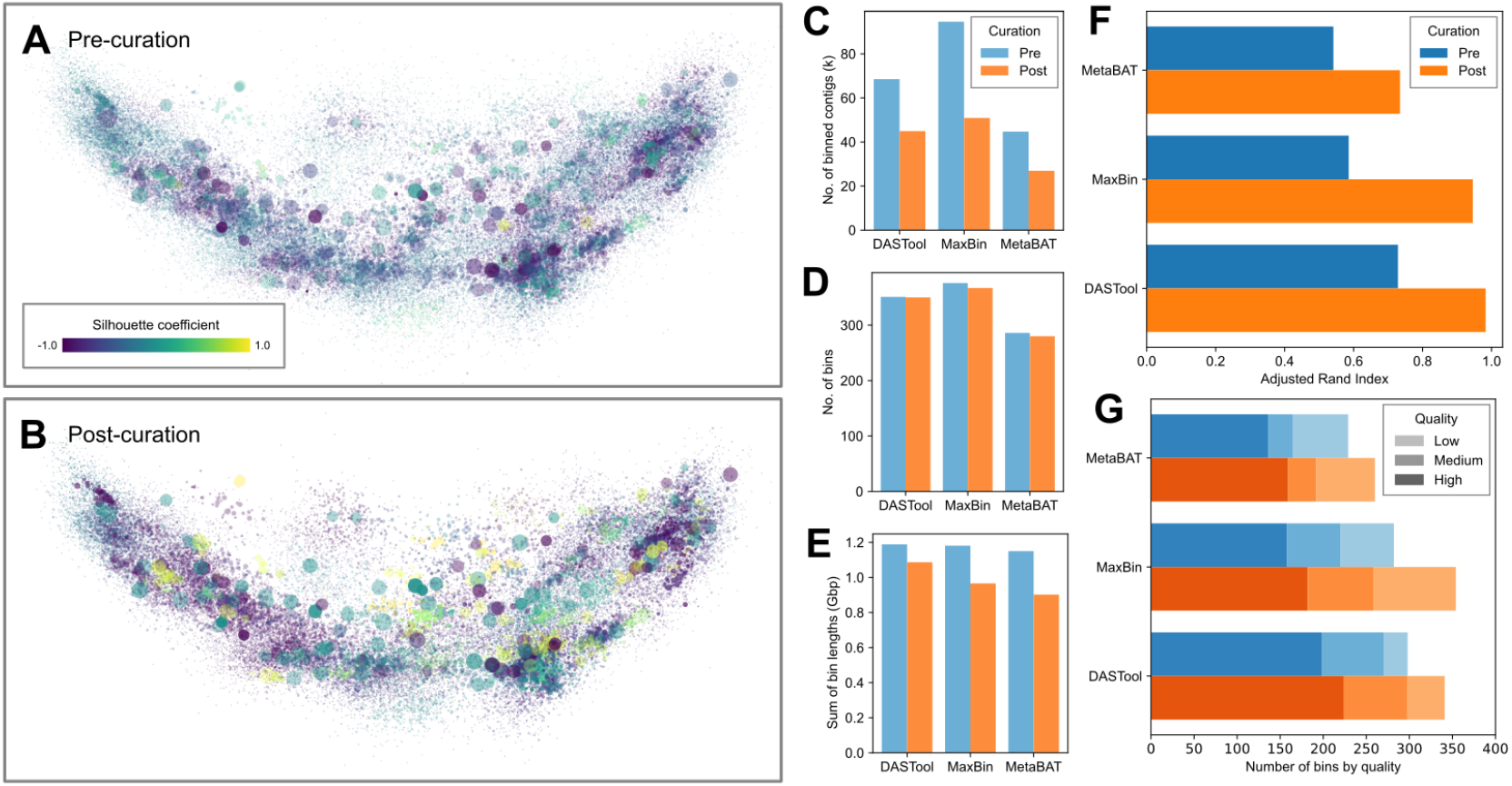
Curation of entire binning plans on the CAMI2 marine dataset. **A**, **B**. The gold standard assembly was visualized using PCA on tetranucleotide frequencies, showing DASTool-binned contigs pre- (**A**) and post- (**B**) curation using BinaRena. Contigs are colored by the silhouette coefficient calculated by BinaRena. Marker size (radius) is proportional to the cube root of contig length. Marker opacity is proportional to the 4th power root of average contig coverage of 10 samples. **C**-**G**. Metrics of three binning plans (generated by DASTool, MaxBin and MetaBAT, respectively) pre- and post-curation. **C**. Total number of contigs in bins. **D**. Total number of bins. **E**. Total length of contigs in bins. **F**. Adjusted Rand index between each binning plan and the ground truth as calculated by BinaRena. **G**. Numbers of high-, medium- and low-quality MAGs, defined following ^1^ based on CheckM-inferred completeness and redundancy (contamination) scores. Specifically, high quality: ≥ 90% complete, < 5% redundant; medium quality: ≥ 50% complete, < 10% redundant; and low quality: < 50% complete, < 10% redundant. Bins that do not match any catalog (i.e., ≥ 10% redundant) are excluded.

## Discussion

We developed BinaRena to support researchers to more effectively and comprehensively visualize and operate on metagenomic datasets. In this work, we have demonstrated that BinaRena can assist human researchers to quickly identify patterns at the community scale with taxonomic and functional relevance in addressing biological questions, while also isolating relevant MAGs from the background. In addition, we have illustrated issues that can arise from solely using automated binners, and that the use of BinaRena can aid in both identification and improvement from above mentioned issues. Even used as a *de novo* binner, BinaRena could yield MAGs with comparable or even better quality than the best result of several automatic binners. Meanwhile, it is effective in curating binning plans computed by automatic binners, and achieving improved quality of the recovered MAGs.

BinaRena’s ease of operation and versatility facilitate metagenomic analysis for both novice and expert users. Being a dependency-free, client-end single webpage, BinaRena is among the easiest of all bioinformatics tools in terms of deployment and use. This characteristic also grants potential for effortless integration of BinaRena into current metagenomics workflows. In contrast to the simplicity in start-up, BinaRena has rich features that permit complex operations on metagenomic data. Meanwhile, the program’s deliberate user interface (UI) design provides an efficient and comfortable workspace for human operators, and this is of importance because the exploration of complex data requires labor and concentration. Noting its high customizability, we envision that BinaRena may also be useful in other research tasks involving classification, clustering, and/or ordination, although further work is needed to establish this point.

While being a useful tool for microbiome researchers, BinaRena is not meant to replace automatic binners. The analysis is highly impacted by human behavior, which could introduce bias. Careful documenting and reasoning (as done in this work) ensure reproducibility of one analysis, but do not warrant generalization of the protocol to other cases. We recommend the adoption of BinaRena in addition to automatic workflows, the results of which are also useful input for BinaRena, as demonstrated above. On top of all, BinaRena is suitable for data overview, hypothesis generation, and sanity check of analysis results. Beyond, BinaRena lets the researcher focus on individual MAGs that are of high relevance to the research topic. Lastly, BinaRena can help if the research goal is to maximize the quality of an entire binning plan, although this would require significant human labor.

The pursuit of decoding complex metagenomic data and deconvoluting them into original organismal entities is of central importance yet so far challenging. BinaRena represents progress in one direction of multiple to the solution of this problem. Future efforts should be attributed to better integration of algorithms and human factors into a semi-supervised workflow that simultaneously achieves high accuracy, interpretability, and reproducibility.

## Materials and Methods

### The Maquia Peatland dataset

The Maquia peatland (MAQ) metagenomes were sampled in the Yanayacu-Maquia Conservation Concession, Peru (6°22′ S 74°53′ W) in October 2015. Six samples were collected from soil cores at three spatial intervals 50m apart at depths of both 10cm and 20cm. DNA extraction was performed using the MicroSoil kit (QIAGEN, CA) following the general protocol proposed by the earth microbiome project ^57^. High-throughput sequencing was performed on an Illumina NovaSeq platform at JGI, NM as part of their 2015 Community Sequencing Program.

Sequencing data were processed using Trimmomatic v0.40 ^58^. Quality-trimmed sequencing data were deposited at JGI for MAQ (Ga0314862-Ga0314867).

### The Travelers’ Diarrhea dataset

The Travelers’ Diarrhea (TD) dataset ^33^ contains 29 metagenomic samples, sequenced from fecal materials of individuals who traveled from the United States to Mexico or India between 2005 and 2010. Twenty-two subjects developed TD but were tested negative for common TD pathogens, implicating the presence of novel pathogens, whereas the remaining seven were healthy. We re-analyzed the published sequencing data (NCBI PRJNA382010) using currently adopted workflows (see below). The metagenomes were assembled separately due to the lack of shared pathogenic profiles. Two samples, #76 and #50076, which were shown to contain closely related pathogens ^33^, were selected for demonstrating BinaRena’s functionality in this study.

### The CAMI2 marine dataset

The 2nd CAMI Challenge ^59^ marine metagenomes (Illumina) gold standard assembly (GSA) was retrieved from PUBLISSO (https://doi.org/10.4126/FRL01-006425521). It contains 10 samples, simulated to represent microbial communities at different seafloor locations of a marine environment. Contigs that are at least 2,000 bp, totaling 159,957 contigs, 1.816 Gbp, were used for binning. The per-sample abundance values were used in this study to assist manual curation of binning plans. The ground truth genome assignments were retrieved from the CAMI GitHub repository (https://github.com/CAMI-challenge/second_challenge_evaluation/tree/masterbinning/genome_binning/marine_dataset/data/ground_truth/gsa_pooled_mapping_short.binning)

### Assembly and automatic binning of metagenomic datasets

Both MAQ and TD metagenomes were co-assembled using MegaHit v1.2.9 ^41^ using the “--meta” preset. Resulting contigs were filtered based on a minimum length of 2,000 bp and an average coverage greater than 1× over 90% of the contig length. Metagenomic reads from each sample were mapped back to contigs using Bowtie2 v2.3.5.1 ^60^ and depth profiles were generated using the “jgi” script provided in MaxBin2 v2.2.7 ^16^. For all four datasets (MAQ, TD, and marine) filtered contigs were binned using MetaBAT2 v2.2.15 ^17^ and MaxBin2 ^16^ with default settings. Results from the these binning plans were consolidated using DASTool v1.1.3 ^18^. Resulting MAGs were assessed for quality using CheckM v1.1.3 ^38^ and GTDB-tk v1.7.0 ^42^ was used to determine taxonomy of all bins. Bins with both completeness and contamination scores equal to zero according to CheckM were not investigated.

### Data preparation for BinaRena

Length, coverage, GC content, and *k*-mer frequencies (*k* = 4, 5, and 6) of individual contigs were calculated using previously published scripts ^16,61^. The *k*-mer frequency profiles were subject to three mainstream dimensionality reduction methods: PCA ^62^, t-SNE ^63^ (implemented in scikit-learn v1.0.2) and UMAP ^64^ (implemented in umap-learn v0.5.3). Prior to the t-SNE analysis, the dataset was processed using PCA to retain 50 dimensions. The Barnes-Hut approximation ^65^ was used to accelerate the t-SNE calculation, following previous works ^36,66,67^. The UMAP analysis was also based on the same 50 PCA-reduced dimensions. Contigs from the MAQ and TD datasets were annotated using Kofamscan v1.3.0 ^68^ against KOfam release 2022-03-01. Taxonomy was assigned to contigs using Kraken2 v2.1.2 ^43^ with default settings against the GTDB release 202 ^69^.

## Code and data availability

The source code of BinaRena is publicly available at: https://github.com/qiyunlab/binarena, under the BSD-3-Clause license. The program is compatible with most modern web browsers (such as Chrome, Firefox, Safari and Edge) and operating systems (such as Windows, MacOS and Linux). Running the program requires no dependency. The data and scripts presented in this manuscript are publicly available at: https://github.com/pavia27/BinaRena-manuscript.

## Acknowledgements

We are grateful to Dr. Sarah Highlander and Dr. Rob Knight for insightful discussions on this study. This work is supported in part by an Arizona State University start-up grant to Q.Z., and NSF DEB project 1749252, JGI CSP No 166 to H.C.-Q..

