## Supplementary Information for "BinaRena: a dedicated interactive platform for human-guided exploration and binning of metagenomes"

Pavia et al.

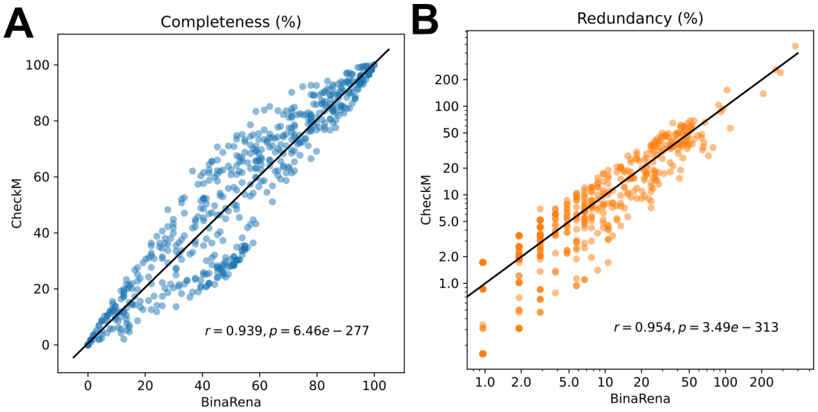


**Figure S1**. Correlation between completeness (**A**) and redundancy (contamination) (**B**) values calculated by BinaRena and CheckM. A total of 596 bins recovered by MaxBin and MetaBAT from the MAQ dataset were evaluated. The CheckM marker gene set for domain Bacteria was used, which contains 104 genes arranged in 58 sets. The regression line (black) is plotted in each panel. The Pearson's correlation coefficient (*r*) and its *p*-value are marked under the plot.

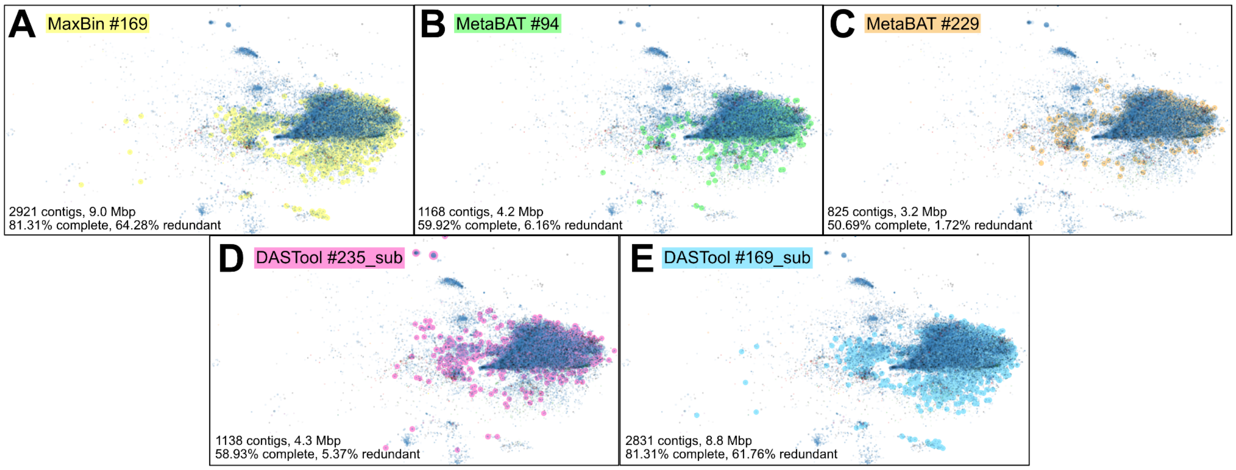


**Figure S2**. *Sulfotelmatobacter* MAGs identified by three automatic binners. Scatter plots were defined by t-SNE on *k*-mer ($k = 6$) frequencies. Marker size (radius) is proportional to contig length. Marker opacity is proportional to the cube root of contig coverage in location 3 at depth of 20cm. **A**. MaxBin's result. **B**, **C**. MetaBAT's results. **D**, **E**. DASTool's results which are both subsets of MaxBin results. **D** came from a bin with high redundancy (40.89%) and only classified to the family level (Koribacteraceae) and **E** is a subset of panel **A**.

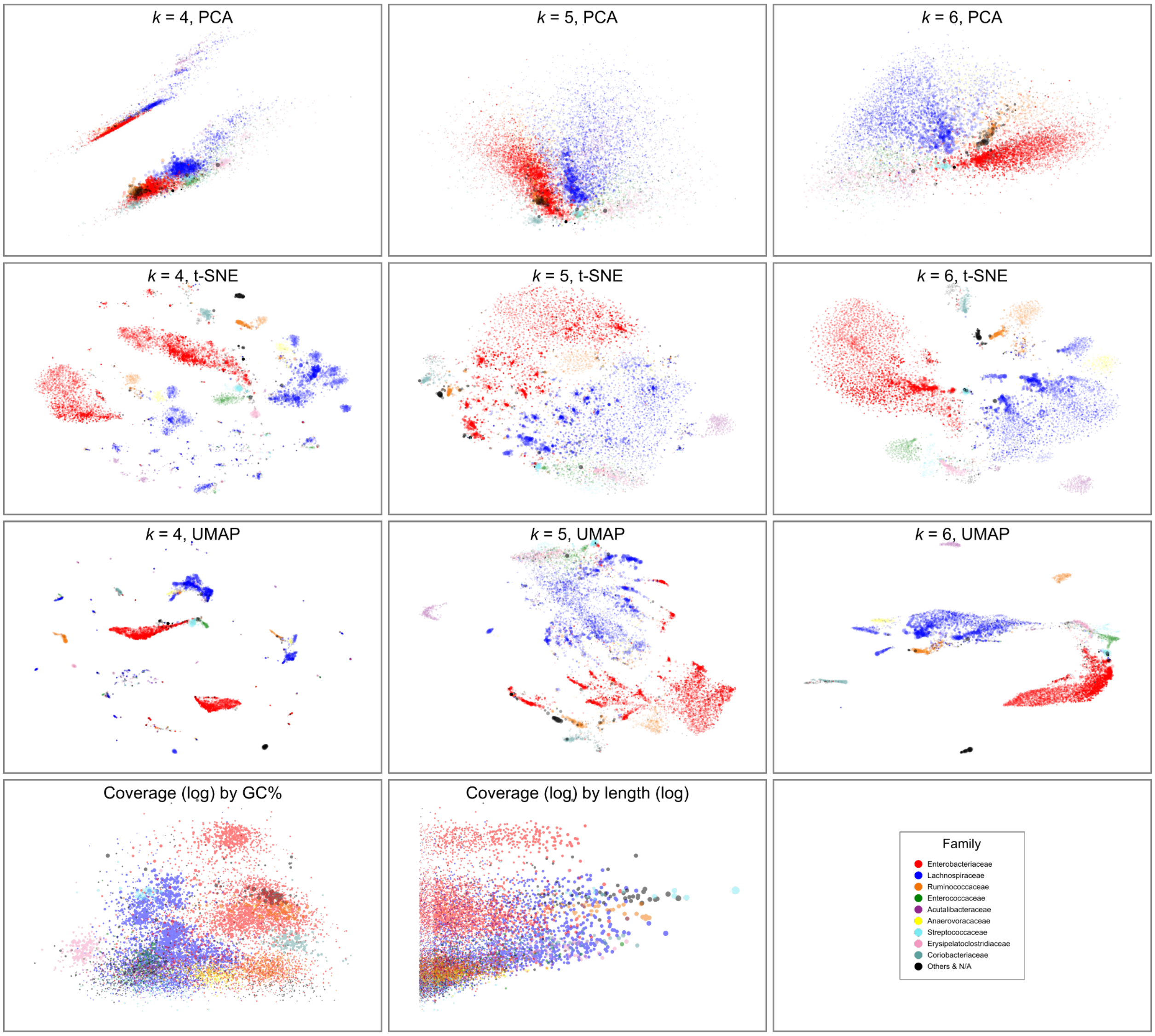


**Figure S3**. Various views of the TD metagenome #76. Three dimensionality reduction methods, PCA, t-SNE, and UMAP, were applied to *k*-mer frequency profiles with $k = 4$, 5, and 6. In addition, the coverage (log) was plotted against GC content and contig length (log). Marker size (radius) is proportional to the cube root of contig length. Except for the last two plots (in which contig coverage is the *y*-axis) , marker opacity is proportional to the square root of contig coverage. Colors are assigned to the top nine most abundant families. The color codes are identical to that of Fig. 3A, B.

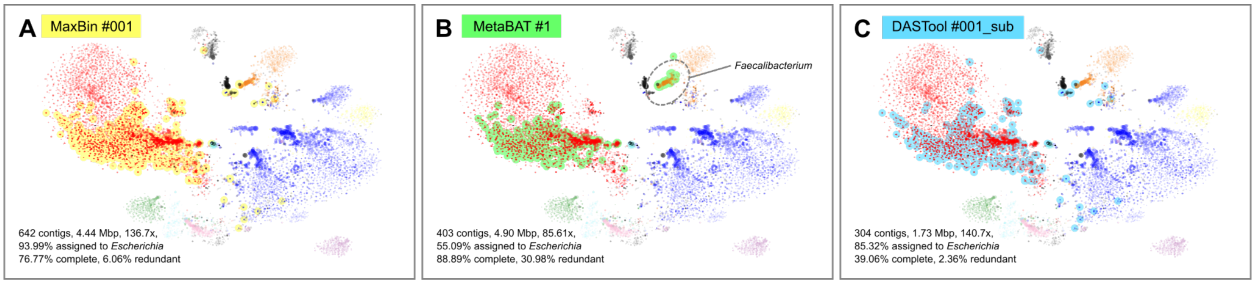


**Figure S4**. Comparison of an *Escherichia* MAG identified by three automatic binners. Scatter plots were defined by t-SNE on *k*-mer ($k = 6$) frequencies. Marker size (radius) is proportional to the cube root of contig length. Marker opacity is proportional to the square root of contig coverage. **A**. MaxBin's result (see also Fig. 3G), which has the highest consistency with the manually identified MAG (Fig. 3F). **B**. MetaBAT's result (also DASTool's primary result), which contains a proportion of the *Escherichia* contigs plus a separate contig cluster assigned to genus *Faecalibacterium* (dashed circle). **C**. DASTool's secondary result (equivalent to the MaxBin bin excluding the MetaBAT bin).

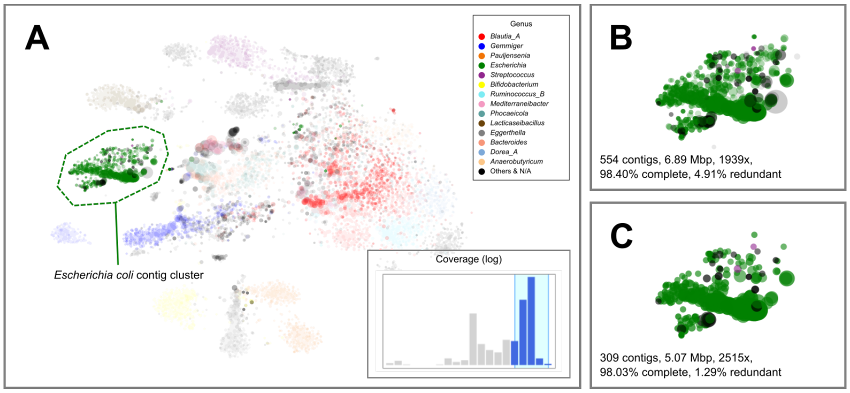


**Figure S5**. Recovery of a pathogenic *Escherichia coli* MAG from TD sample #50076 (19.82 Gbp raw reads, 9,816 contigs totaling 62.2 Mbp), which contains multiple *E. coli* strains. Scatter plot was defined by t-SNE on *k*-mer ($k = 6$) frequencies. Marker size (radius) is proportional to the cube root of contig length. Marker opacity is proportional to the cube root of contig coverage. Colors were assigned to the top 14 most abundant genera in the sample. **A**. View of the entire assembly. A cluster of contigs mainly assigned to *Escherichia* was selected (dashed polygon). **B**. The selected cluster of contigs. Its coverage profile exhibits a multi-modal pattern (inset of A). Therefore, the top five out of 20 bins of the histogram were retained. **C**. The retained contigs, which represent a putative *E. coli* MAG.

**Table S1**. Comparison of functionality of BinaRena and Elviz.

| Feature | BinaRena | Elviz | Feature | BinaRena | Elviz |
|---|---|---|---|---|---|
| Deployment methods | Local or remote (no installation or server required) | Web server (integrated into JGI) | Contig searching | Search by field and data type (keyword and data range) | Simple keyword searching |
| Visualization method | Scatter plot | Scatter plot | Contig annotation | Yes (feature set) | Yes (graphic, with genome map) |
| Customizable plot aesthetics | *x*- and *y*-axes, size, color (continuous and discrete, multiple palettes), opacity | *x*- and *y*-axes, size, color (discrete only, preloaded) | Contig-to-bin assignment | Add, delete | Reassign |
| | | | Bin management | Add, delete, merge, rename | Add |
| Data transformation methods | Linear, logarithmic, exponential, square / cube / 4th power (root), ranking, logit, arcsine | Linear, logarithmic | Interactive histogram | Yes | No |
| | | | Binning plan evaluation | Yes (silhouette coefficient) | No |
| Data range specifying | Yes | Yes | Binning plan comparison | Yes (adjusted Rand index) | No |
| Contig selection - single | Yes (mouse click) | Yes (mouse click) | Completeness / redundancy calculation | Yes | No |
| Contig selection - group | Yes (polygon) | Yes (lasso) | *k*-mer signature calculation | Yes | No |
| Contig information - single | Yes (side panel) | Yes (floating window) | Dimensionality reduction | Yes | No |
| Contig information - group | Yes (side panel) | No | Data table view | Yes | No |
| Contig masking | Yes | No | Data export | Yes | Yes |
| Contig highlighting | Yes | No | Image export | PNG and SVG | No |
| Contig focusing | Yes | Yes | Link to JGI project | No | Yes |

**Table S2**. Metrics of contig clusters / bins in TD sample #76 calculated by BinaRena.

| Fig. | Description | # Contigs | Length (bp) | Coverage (x) | GC (%) | Family | Genus | % Complete | % Redundant |
|---|---|---|---|---|---|---|---|---|---|
| 3C | *Enterobacteriaceae* cluster | 4293 | 19944053 | 47.624 | 52.558 | *Enterobacteriaceae* (98.42%) | ambiguous | 84.85 | 137.37 |
| 3E | *Escherichia* bin post coverage filtering | 644 | 4467795 | 136.386 | 50.517 | *Enterobacteriaceae* (98.8%) | *Escherichia* (95.87%) | 78.11 | 6.4 |
| 3F | *Escherichia* bin post taxonomy filtering | 611 | 4295399 | 137.463 | 50.61 | *Enterobacteriaceae* (99.89%) | *Escherichia* (99.72%) | 75.42 | 3.7 |
| 3G, S4A | MaxBin #001 | 642 | 4444443 | 136.713 | 50.748 | *Enterobacteriaceae* (96.77%) | *Escherichia* (93.99%) | 76.77 | 6.06 |
| 3H | MaxBin #001 post curation | 607 | 4230607 | 138.208 | 50.806 | *Enterobacteriaceae* (98.91%) | *Escherichia* (98.74%) | 74.07 | 3.03 |
| S4B | MetaBAT #1 | 403 | 4896109 | 85.608 | 54.482 | *Enterobacteriaceae* (55.41%) | *Escherichia* (55.09%) | 88.89 | 30.98 |
| S4C | DASTool #001_sub | 304 | 1734512 | 140.682 | 50.037 | *Enterobacteriaceae* (91.85%) | *Escherichia* (85.32%) | 39.06 | 2.36 |

**Table S3**. Metrics of contig clusters / bins in TD sample #50076 calculated by BinaRena.

| Description | # Contigs | Length (bp) | Coverage (x) | GC (%) | Genus | % Complete | % Redundant |
|---|---|---|---|---|---|---|---|
| MaxBin #001 | 83 | 1667686 | 3075.709 | 51.049 | *Escherichia* (99.15%) | 40.97 | 0.37 |
| MaxBin #002 | 84 | 1592021 | 2500.841 | 51.304 | *Escherichia* (99.22%) | 36 | 0.18 |
| MaxBin #003 | 68 | 1202044 | 2111.743 | 49.921 | *Escherichia* (98.95%) | 18.37 | 0.12 |
| MetaBAT #9 | 89 | 1898150 | 3007.073 | 51.429 | *Escherichia* (99.25%) | 51.54 | 0.37 |
| MetaBAT #28 | 99 | 2249941 | 2269.656 | 50.745 | *Escherichia* (100.00%) | 43.24 | 0.31 |
| Blob of *E. coli* contigs | 554 | 6891371 | 1938.569 | 49.515 | *Escherichia* (89.25%) | 98.4 | 4.91 |
| Top 5 bins out of 20 | 309 | 5069114 | 2515.359 | 50.492 | *Escherichia* (96.61%) | 98.03 | 1.29 |
